# Allele-specific CRISPR/Cas9 genome editing of the single-base P23H mutation for rhodopsin associated dominant retinitis pigmentosa

**DOI:** 10.1101/197962

**Authors:** Pingjuan Li, Benjamin P. Kleinstiver, Mihoko Y. Leon, Michelle S. Prew, Daniel Navarro-Gomez, Scott H. Greenwald, Eric A. Pierce, J. Keith Joung, Qin Liu

## Abstract

Treatment strategies for dominantly inherited disorders typically involve silencing or ablating the pathogenic allele. CRISPR/Cas nucleases have shown promise in allele-specific knockout approaches when the dominant allele creates unique protospacer adjacent motifs (PAMs) that can lead to allele restricted targeting. Here, we present a spacer-mediated allele-specific knockout approach that utilizes both SpCas9 variants and truncated single guide RNAs (tru-sgRNAs) to achieve efficient discrimination of a single-nucleotide mutation in rhodopsin (*Rho*)-P23H mice, a model of dominant retinitis pigmentosa (RP). We found that approximately 45% of the mutant P23H allele was edited at DNA level, and that the relative RNA expression of wild-type *Rho* was about 2.8 times more than that of mutant *Rho* in treated retinas. Furthermore, the progression of photoreceptor cell degeneration in outer nuclear layer was significantly delayed in treated regions of the *Rho*-P23H retinas at five weeks of age. Our proof-of-concept study therefore outlines a general strategy that could potentially be expanded to examine the therapeutic benefit of allele-specific gene editing approach to treat human P23H patient. Our study also extends allele-specific editing strategies beyond discrimination within the PAM sites, with potentially broad applicability to other dominant diseases.

## Introduction

Conventional gene augmentation therapies have shown great promise in treating recessive diseases, such as inherited retinal degenerations (IRDs) and hemophilia,^1, 2^ yet their application to dominant diseases is limited due to the requirement for silencing or ablating the gain-of-function or dominant-negative mutant alleles. While alternative RNA-based suppression-replacement using ribozyme, zinc-finger-based artificial transcription factors, or RNAi-based approaches to transiently silence or degrade both mutated and normal RNA transcripts, in combination with supplementation of an exogenous wild-type transgene have been actively pursued, the allelic specificity, longevity of the therapy, and regulation of gene expression levels have posed significant challenges for broad application.^3-6^ Recently developed CRISPR/Cas9 gene editing technologies present an alternative approach to treat dominant diseases by addressing these challenges.

The CRISPR/Cas system has been revolutionary for genome-editing because the Cas9 nuclease can be programmed to generate double-strand DNA breaks at specified genomic sites when targeted by a single guide RNA (sgRNA),^7^ which dictates much of the target site specificity.^8^ The Cas9/sgRNA complex scans the genome for the presence of a protospacer adjacent motif (PAM), the critical first step of DNA target site recognition.^8^ Recent studies in mice have shown that when the disease-causing allele harbors unique PAM sequences that are not present in its wild-type counterpart, allele-specific genome editing by selective targeting and permanent inactivation of the mutant allele could be achieved while leaving the wild-type allele functionally intact.^9-11^ However, many dominant alleles do not carry sequences that generate unique PAM sites, making them refractory to this type of allele-specific editing strategy. One potential approach to expand the applicability of allele-specific genome editing is to develop strategies in which the disease-causing allele can be discriminated from the wild-type allele when the mutations are within the spacer region of the Cas9/sgRNA target site. It is a widely-recognized challenge that Streptococcus pyogenes Cas9 (SpCas9) is generally ineffective at distinguishing between single nucleotide mismatches in much of its spacer sequence,^12-15^ however, improvements to SpCas9 targeting specificity by truncating the sgRNA may be leveraged to overcome this hurdle.^16^ Furthermore, recent studies in human iPSCs and rat embryos have provided some encouraging evidence supporting the use of mismatches in the spacer sequence to achieve allele-specific genome editing,^17-19^ including a most recent report of allele-specific editing against a human *Rhodopsin*-P23H allele in hiPSCs.^20^ In the present study, we investigated the feasibility of using this spacer-mediated allele-specific CRISPR/Cas9 gene editing to target an endogenous single-base missense mutation at its native locus in a *Rhodopsin*-P23H knockin mouse model that genetically and phenotypically recapitulates human autosomal dominant retinitis pigmentosa (adRP).^21, 22^

Retinitis pigmentosa (RP) is the most common inherited retinal degeneration (IRD), affecting 1 in 3500 people.^23^ Vision loss in IRDs is caused by progressive photoreceptor cell dysfunction and death.^23^ Mutations in 23 genes have been reported to cause adRP,^24^ including over 180 mutations in the rhodopsin (*RHO*) gene, which accounts for over 25% of adRP cases.^25, 26^ Approximately half of the *RHO*-associated adRP cases are caused by the P23H mutation.^25, 26^ The mutant P23H rhodopsin protein is thought to misfold and co-aggregate with wild-type rhodopsin, resulting in a gain-of-function or dominant negative effect in rod photoreceptor cells.^3, 6^ In this study, by designing and testing combinations of sgRNAs and Cas9 variants, we have demonstrated effective and specific knockout of the mutant P23H allele in the retina of *Rho*-P23H heterozygous mice. In treated mice we observed a significant decrease of the expression of mutant P23H transcript, which in turn led to preservation of the thickness of photoreceptor cell layer in the treated region of the retina. The proof-of-concept presented in this study supports the notion that the spacer-mediated allele-specific CRISPR/Cas9 gene editing approach may be applicable not only to *RHO*-P23H associated adRP but also to other dominant disorders.

## Materials and Methods

### Mouse Husbandry

Our study conforms to the Association for Research in Vision and Ophthalmology Statement for the Use of Animals in Ophthalmic and Vision Research. All procedures were approved by the Animal Care and Use Committee of the Massachusetts Eye and Ear Infirmary. The *Rho*-P23H mouse line was purchased from the Jackson Lab (stock No. 017628) and maintained on a 12h: 12h light/dark cycle. F1 heterozygous progeny generated by crossing P23H homozygous mice with wild-type mice, and F2 progeny of wild-type, heterozygous and homozygous mice were generated by intercrossing F1 heterozygous mice.

### Plasmid constructs

Plasmids encoding pCAG-SpCas9-VQR-P2A-EGFP, pCAG-SpCas9-VRQR-P2A-EGFP, and pCAG-KKH-SaCas9-P2A-EGFP sequences were generated via isothermal assembly. The Cas9 expression constructs also encode an *EGFP* gene separated from the nuclease by a P2A sequence to facilitate the identification and sorting of transfected cells. Oligonucleotides to clone sgRNAs were synthesized by Integrated DNA Technologies (Coralville, Iowa, USA), annealed, and ligated into BsmBI digested BPK2660 (Addgene #70709) or BPK1520 (Addgene #65777) for SaCas9 and SpCas9 sgRNAs, respectively.

### Subretinal injection

P0-P2 pups were anesthetized by hypothermia and plasmid DNAs were injected into subretinal space using standard methods.^27^ Briefly, 0.5μl of DNA solution containing Cas9-encoding plasmid with or without the sgRNA bearing plasmid (absolute amount of DNA is 0.6 μg for Cas9 and 0.3μg for sgRNA, respectively) was injected using a 34G Hamilton syringe, followed by *in vivo* electroporation using five 90V, 50ms square pulses that were delivered with an Electro Square Porator (BTX, Holliston, MA).

### Retina dissociation and cell sorting

Eyes were enucleated at P5-P7 days for DNA extraction or at P14 days for RNA isolation, respectively. Retinas were isolated in BGJB medium (Thermo Fisher Scientific, Waltham, MA) on ice under a fluorescent dissection microscope to document the transfected region, and then dissociated into single cells by incubation in Solution A containing 1mg/ml pronase (Sigma-Aldrich, St. Louis, MO) and 2mM EGTA in BGJB medium at 37°C for 20 minutes. Solution A was gently removed, followed by adding equal amount of Solution B containing 100 U/ml DNase I (New England Biolabs, Cambridge, MA), 0.5% BSA, 2mM EGTA in BGJB medium. Cells were collected and re-suspended in 1 X PBS, filtered through a Cell Strainer (BD Biosciences, San Jose, CA), and submitted for fluorescence activated cell sorting (FACS). EGFP positive and negative cells were collected using a Cytomation MoFlo Cell Sorter (Cytomation, Fort Collins, CO).

### Genomic DNA extraction and PCR

Genomic DNA was extracted from sorted cells using DNA Extraction solution (Epicenter, Madison, WI), and 100ng of genomic DNA was used as PCR template to amplify the sequence flanking the sgRNAs on-target and predicted top 10 off-target sequences. Primer sets are listed in **Supplementary Table 1**. The PCR amplicons were then subjected for Next Generation Sequencing (NGS) analysis.

### RNA extraction and RT-PCR

Total RNA from sorted EGFP positive and negative cells at P14 was extracted using the RNeasy Micro Kit (Qiagen, Hilden, Germany). Total RNA was reverse transcribed with the SuperScript IV First-Strand Synthesis System (Thermo Fisher Scientific, Rockford, IL). cDNA was PCR amplified using the condition described above and subjected for NGS analysis.

### Targeted deep sequencing

The on-target and off-target activities of Cas9-sgRNA pairs used in this study were evaluated by targeted deep sequencing. Briefly, PCR or RT-PCR amplicons of the target site or predicted off-target sites from treated and untreated cells were analyzed by NGS using the primers listed in **Supplementary Table 1**. Paired-end sequencing of the resulting TruSeq-compatible paired-end Illumina libraries was performed on the Illumina MiSeq platform. Sequencing data was analyzed using the CRISPResso software followed by a custom program that corrects nonsense single nucleotide polymorphisms (SNPs) due to NGS errors and consolidates indel counts^28^. Allele frequencies from treated retinas and untreated heterozygous control retinas were analyzed via calculating the percentage of paired reads for each of the wild-type, P23H, or edited alleles.

### Immunostaining and microscopy

Eyes from Cas9/sgRNA treated mice were isolated and processed for retinal immunostaining experiments five weeks post-injection as described previously.^29^ Retinal cryosections were cut and evaluated using anti-rhodopsin antibody at 1:1000 dilution (MAB5356, EMD Millipore, Billerica, MA), followed by Alexa 555-conjugated secondary antibodies. Images were taken using the Eclipse Ti fluorescence microscope. Outer nuclear layer (ONL) thickness of the EGFP(+) region and the corresponding region in the control retinas were measured.

### Statistical analysis

All data was presented as mean ± SD. The two-tailed unpaired t-test was used for data analysis. Statistical significance was considered when *P* value is < 0.05.

## Results

### Design and allele-specificity screening of sgRNAs for targeting the *Rho*-P23H allele in mouse retina

The endogenous sequence surrounding the mouse *Rho*-P23H mutation does not contain any canonical NGG PAM sites for wild-type SpCas9, so for an alternative approach we designed two sgRNAs (sgRNA1 and sgRNA2) against sites with non-canonical NNCAGT and NGA PAMs that are accessible when using engineered SaCas9-KKH and SpCas9-VQR nucleases, respectively **(Fig. 1a)**.^30, 31^ For these two target sites, the P23H mutation is located either 12 or 4 base pairs (bp) upstream of the PAM in the sgRNA1 or sgRNA2 target sites, respectively. To determine the cleavage efficiency and allele-specificity of the sgRNAs along with their compatible Cas9 variants, DNA plasmids were transfected into the retinas of wild-type, heterozygous and homozygous P23H mice at age P0-P2 via sub-retinal injection and *in vivo* electroporation, followed by NGS analysis of the genomic DNA at 5-7 days post injection. The transfection efficiency was approximately 1-3% on average, measured by the percentage of EGFP positive cells compared to all retinal cells. Targeted deep sequencing analysis of PCR products amplified from the Cas9-gRNA transfected cells revealed robust cutting efficiency, but that neither nuclease-sgRNA pair was able to distinguish the mutant P23H allele from the wild-type *Rho* allele **(Fig. 1b, 1c)**. Cutting efficiencies of 37.8% and 40% were observed for the SaCas9-KKH/sgRNA1 and SpCas9-VQR/sgRNA2 pairs, respectively, in the injected wild-type mice even though both nucleases preferentially targeted the mutant allele in heterozygous mice **(Fig. 1b, 1c, Supplementary Table 2)**. As expected, the frequencies of wild-type and P23H alleles in the control heterozygous retinas were approximately 50% each when evaluated by NGS analysis **(Fig. 1b-1e)**.

**Figure 1.**
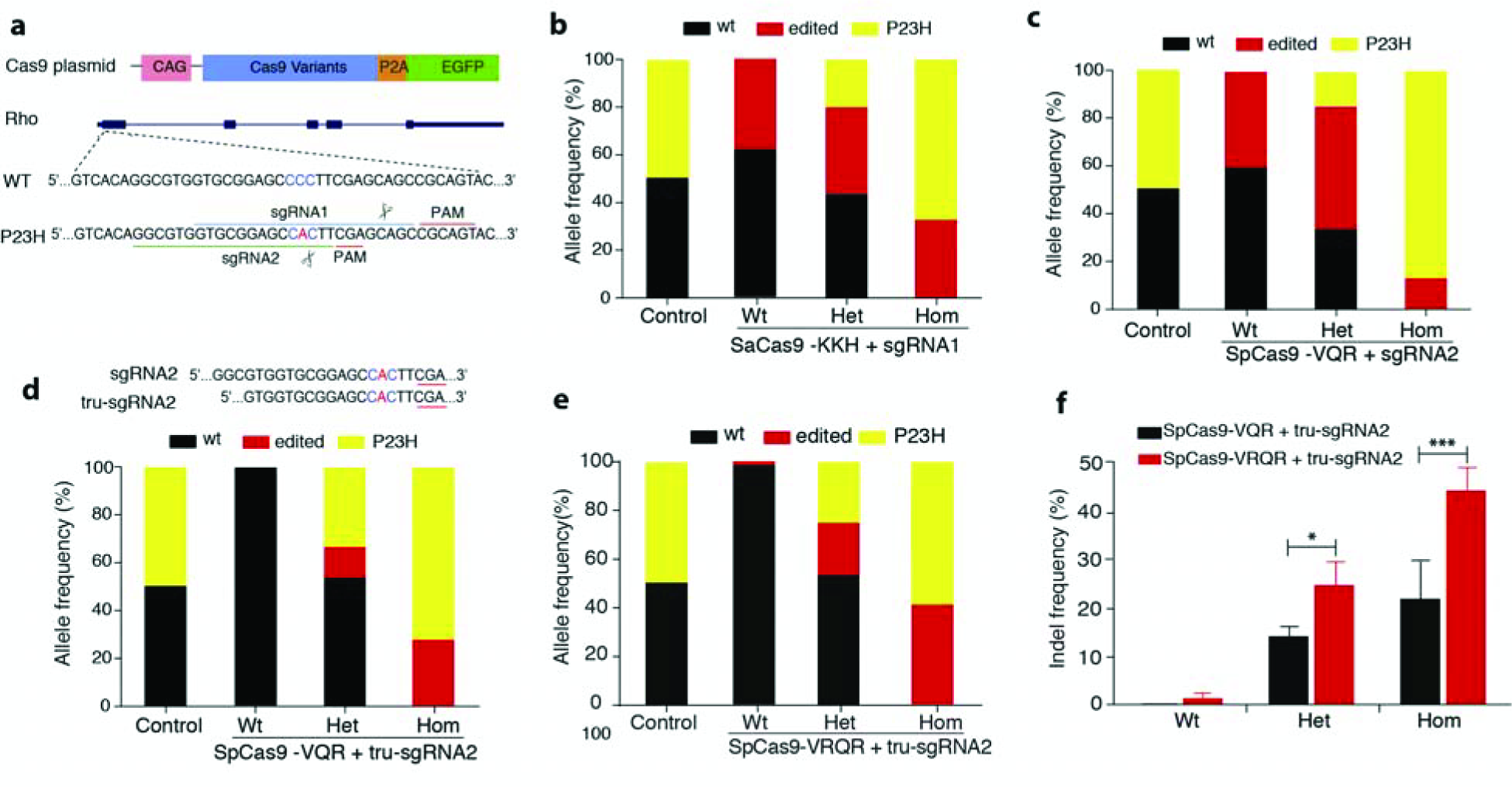
P23H allele-specific sgRNA screening *in vivo*. **(a)** Schematic overview of CRIPSR/Cas9 constructs and their target sites in the *Rho*-P23H allele. Cas9 construct (top panel). PAM sequence is underlined. **(b-d)** Allele frequencies of wt, P23H and edited alleles in injected wild type (wt), P23H heterozygous (het) homozygous (hom) mice and uninjected het mice (control). Allele frequencies were determined by the percentage of paired reads detected by NGS. **(b)** sgRNA1 with SaCas9-KKH; **(c)** sgRNA2 with SpCas9-VQR; **(d)** tru-sgRNA2 + SpCas9-VQR; **(e)** tru-sgRNA2 + SpCas9-VRQR. **(f)** Comparison of targeting efficiency of SpCas9-VQR vs. SpCas9-VRQR combined with tru-sgRNA2 in wt, het and hom mice determined by NGS. * *P* < 0.05; *** P < 0.001 (Student’s T-test, n=3).

Given that both conventional full-length sgRNA1 and sgRNA2 were not capable of allele-specific editing, we therefore examined whether a truncated 17 nt version of sgRNA2(tru-sgRNA2) bearing a shortened 5’ end, an approach shown to increase specificity in a previous study,^16^ could improve allele discrimination. When paired with tru-sgRNA2, SpCas9-VQR exclusively edited the P23H mutant allele with a cleavage efficiency of 28% in the homozygous mutant retinas, and no detectable cleavage in the wild-type controls **(Fig. 1d, Supplementary Table 2)**.

### Improvement of cleavage efficiency of the P23H allele *in vivo*

To potentially improve on-target P23H allele editing efficiency, we next tested tru-sgRNA2 with SpCas9-VRQR, an improved version of SpCas9-VQR that has enhanced activity against NGA PAM sites **(Fig. 1e)**.^32^ On-target editing of the P23H allele with SpCas9-VRQR and tru-sgRNA2 (44.8 ± 4.8%) was significantly improved relative to that of SpCas9-VQR (22.1 ± 8.1%, p<0.001) in homozygous mice, with a very low level of nuclease-induced editing of the wild-type allele (1.3 ± 0.3%) in wild-type mice **(Fig. 1e, 1f)**. These data demonstrate that tru-sgRNA2 paired with SpCas9-VRQR cleaves the P23H allele with greater efficiency, thus this nuclease/guide pair was used in subsequent experiments.

### Indel profiles analysis of the edited P23H allele

To assess whether the insertion or deletion mutations (indels) created by the activity of SpCas9-VRQR and tru-sgRNA2 could lead to the knockout of mutant P23H allele, we analyzed the indel profiles of the edited P23H alleles across 10 independent groups of mice that were treated on different days. We observed that 89.5 ± 3.3% of the indels at the targeted P23H mutation site were frame-shifts **(Fig. 2b, 2c, Supplementary Fig. 1)**. These frame-shift indels are expected to create a premature stop codon at protein position 81aa when the indel was 3n+1bp insertion or –(3n+2)bp deletion or position 142aa when the indel was 3n+2bp insertion or –(3n+1)bp deletion, respectively. The frame-shift indels are predicted to result in nonsense-mediated decay of the edited P23H transcripts.^33^ Moreover, consistent with a previous report, the indel pattern observed was non-random,^34^ as the top two most frequent indels across 10 different experimental replicates were a 1bp insertion (C or A) and 2bp deletion (-CT), accounting for 50.2 ± 4.7% of all indels **(Fig. 2a, 2c, 2d)**.

**Figure 2.**
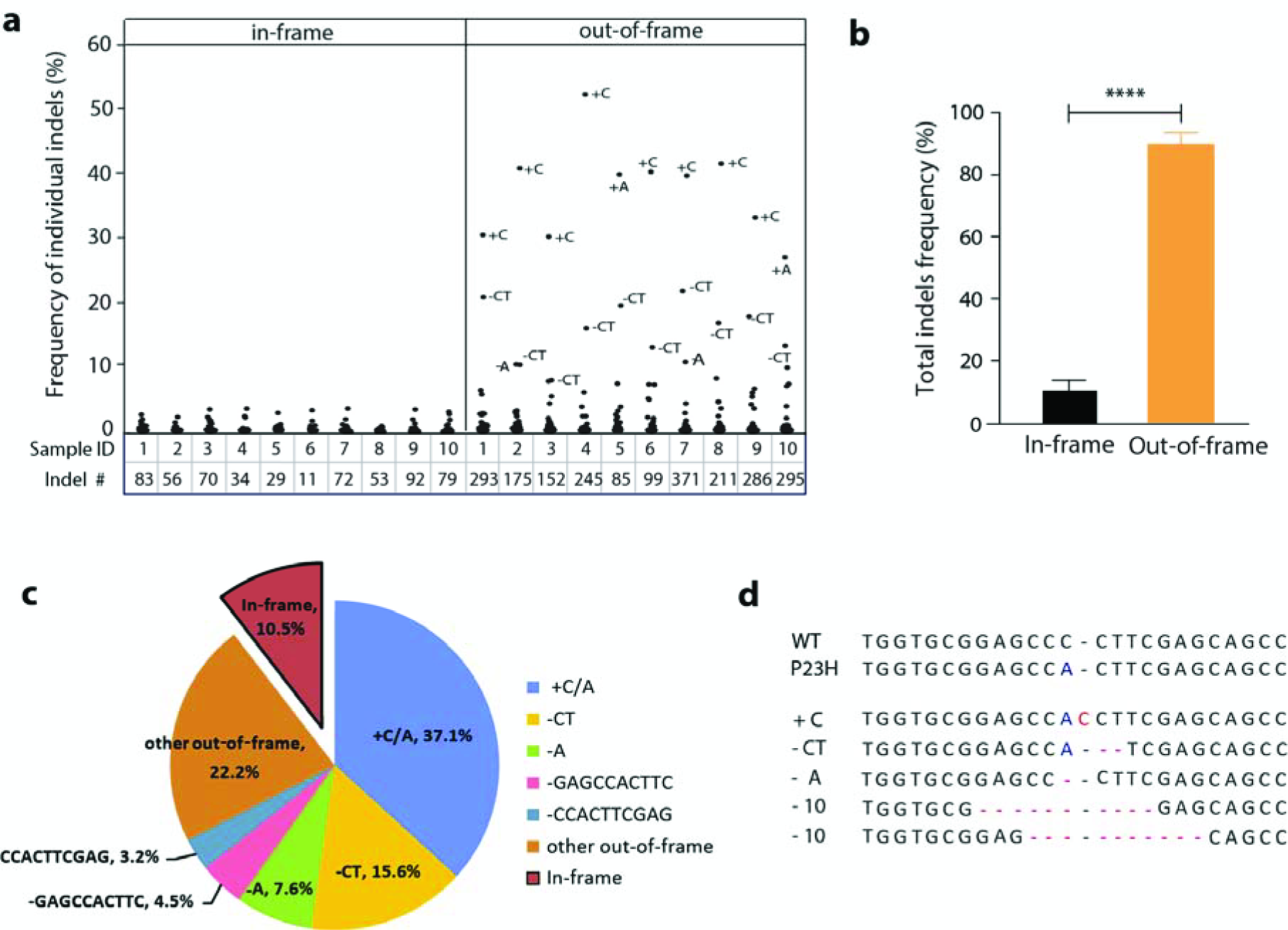
Indel profile of tru-sgRNA2 with VRQR-SpCas9. **(a)** Distribution and frequency of individual in-frame (left) and out-of-frame (right) indels from P23H het mice injected with SpCas9-VRQR and tru-sgRNA2 in 10 independent groups of mice. The total number of indel patterns for each sample is listed at bottom. **(b)** Average percentage of in-frame vs. out-of-frame indels. *** *P* < 0.001 (Student’s T-test, n=10). **(c)** Average percentage of top 5 and other out-of-frame and in-frame indels. **(d)** Representative sequences of top five indel patterns.

### Decreased expression level of P23H mutant mRNA in CRISPR/Cas9 treated cells

We next determined whether the indels introduced by SpCas9-VRQR/tru-sgRNA2 at the P23H DNA level could influence the expression level of mutant P23H mRNA via non-sense mediated decay. We performed targeted deep sequencing of RT-PCR products to compare the amount of mRNA transcripts of wild-type, P23H and edited alleles in heterozygous retinas 14 days after transfection, at which time the expression of rhodopsin is easily detectable. At the DNA level, the percentage of reads for wild-type, P23H and edited alleles were 51.2 ± 2.4%, 25.7 ± 5.0% and 22.8 ± 2.9%, respectively, in the treated cells **(Fig. 3a)**. A comparison at the relative mRNA levels demonstrated that the percentage of wild-type transcript was significantly higher in treated cells versus untreated cells (75.5 ± 2.3% vs. 57.6 ± 0.9%) **(Fig. 3b)**, and the percentage of P23H transcript was significantly lower in the treated cells versus the control samples (22.1 ± 5.8% vs. 42.1 ± 0.8%) **(Fig. 3b)**. Notably, the relative percentage of edited allele was reduced from 22.8% ± 2.9% at DNA level to only 4.8% ± 4.2% at mRNA level in the treated cells **(Fig. 3b)**. Sequence analysis also revealed that the majority of the edited alleles detected at mRNA levels were in-frame indels (data not shown). Thus, the ratio of wild-type mRNA vs. mutant mRNA was significantly increased in the treated cells (2.8 ± 0.35) compared to that in the control cells (1.37 ± 0.05) **(Fig. 3c)**. When mRNA levels were normalized to the wild-type allele, the ratio of wt vs. mutant mRNA was 1: 0.35 in the treated cells compared to 1: 0.73 in the untreated cells **(Fig. 3d)**, resulting a 38% decrease of the mutant mRNA in the treated cells **(Fig. 3e)**. This result is consistent with our finding at DNA level that 89.5% of edited alleles harbor an out-of-frame indel. As the majority of the aberrant transcripts of mutant alleles with out-of-frame indels are predicted to be removed via nonsense mediated decay, this suggests that of all cells transfected with SpCas9-VRQR/tru-sgRNA2, presumably 38% of them express only wild-type rhodopsin mRNA.

**Figure 3.**
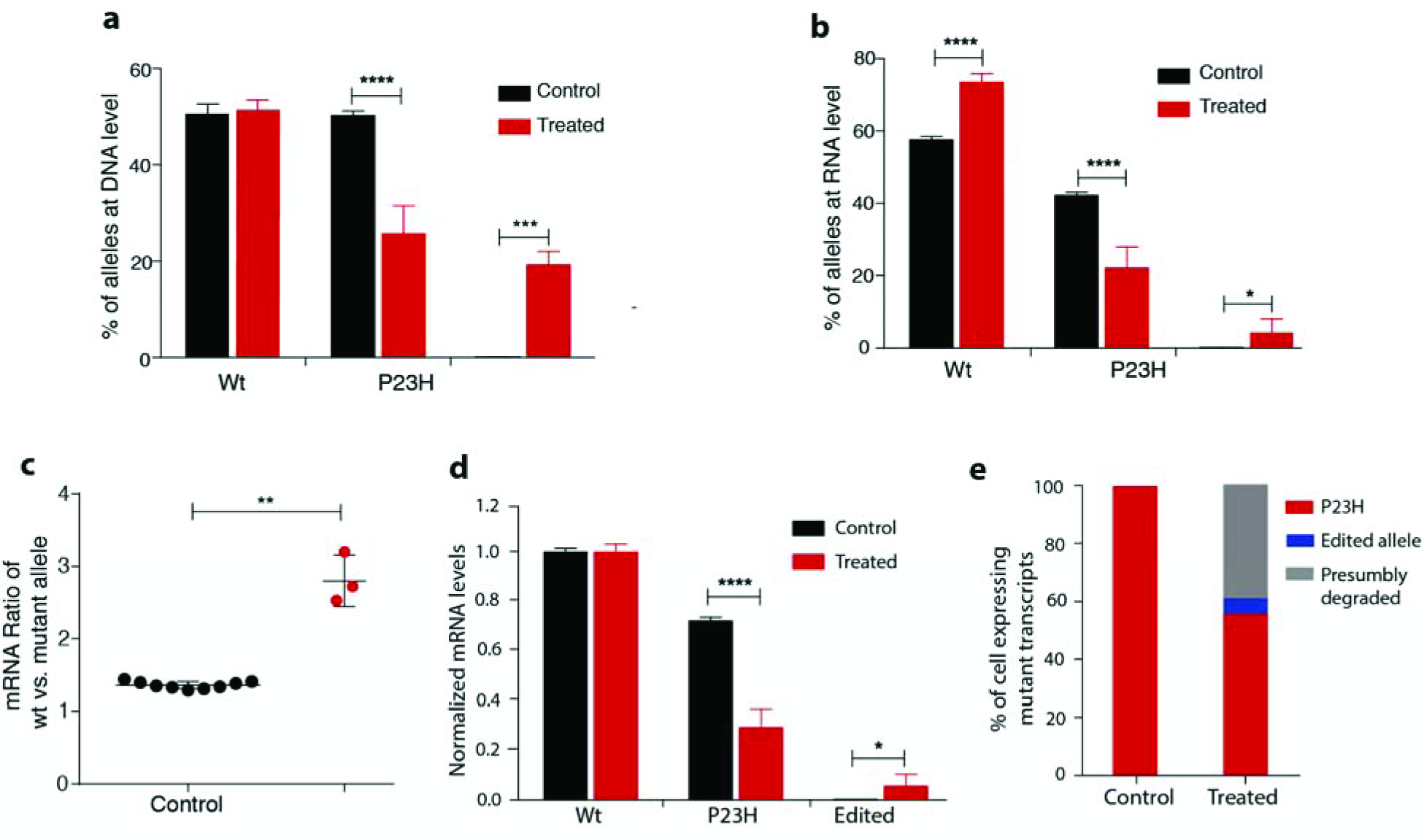
Expression of P23H allele after gene editing with tru-sgRNA2 and SpCas9-VRQR in P23H heterozygous mice. Relative DNA level **(a)** and RNA level **(b)** of wt, P23H allele and edited allele in the treated retinas (n=3) with tru-sgRNA2 and SpCas9-VRQR vs. untreated retinas (n=9) in P23H het mice 14 days post-injection. **(c)** The ratio of wt to mutant (P23H + Edited) alleles at mRNA level in the treated (n=3) and untreated retinas (n=9). **(d)** Normalized mRNA level of Wt, P23H, and indels in the treated (n=3) and untreated retinas (n=9). **(e)** Percentage of cells presumably expressing Wt/P23H, Wt/Edited P23H, and Wt only in the treated and untreated retinas. * *P* < 0.05; ** *P* < 0.01; *** *P* < 0.001; **** *P* < 0.0001.

### Photoreceptor cell preservation in treated *Rho*-P23H mouse retinas

In order to gauge whether ablation of the mutant P23H allele could result in prevention or delay of photoreceptor degeneration, we evaluated the retinal structure in the regions transfected with SpCas9-VRQR/tru-sgRNA2 plasmids in *Rho*-P23H heterozygous mice at age of 5 weeks, at which time the retinas of P23H mice demonstrate significant degeneration histologically and functionally.^21, 22^ Immunofluorescence analysis showed that at the transfected region of retina, approximately 30% of photoreceptor cells were treated with Cas9 components, as counted by the EFGP positive cells over all cells stained with DAPI in outer nuclear layer (ONL) **(Fig. 4a)**. Measurement of ONL thickness revealed that there were significantly more photoreceptor cells in the treated EGFP positive region (five to six rows of cells, 37.3 ± 3.7 μm) than the adjacent untreated EGFP negative area (three to four rows of cells, 25.9 ± 3.9 μm; P < 0.05) **(Fig. 4a, 4b)**. Also, we observed that the rhodopsin signal in the ONL was less intense in the GFP-positive region than that in the GFP-negative region **(Fig. 4a, left panels, insets)**. These results suggest that CRISPR/Cas9-mediated disruption of the P23H mutant *RHO* allele resulted in photoreceptor cell preservation in the treated retinas.

**Figure 4.**
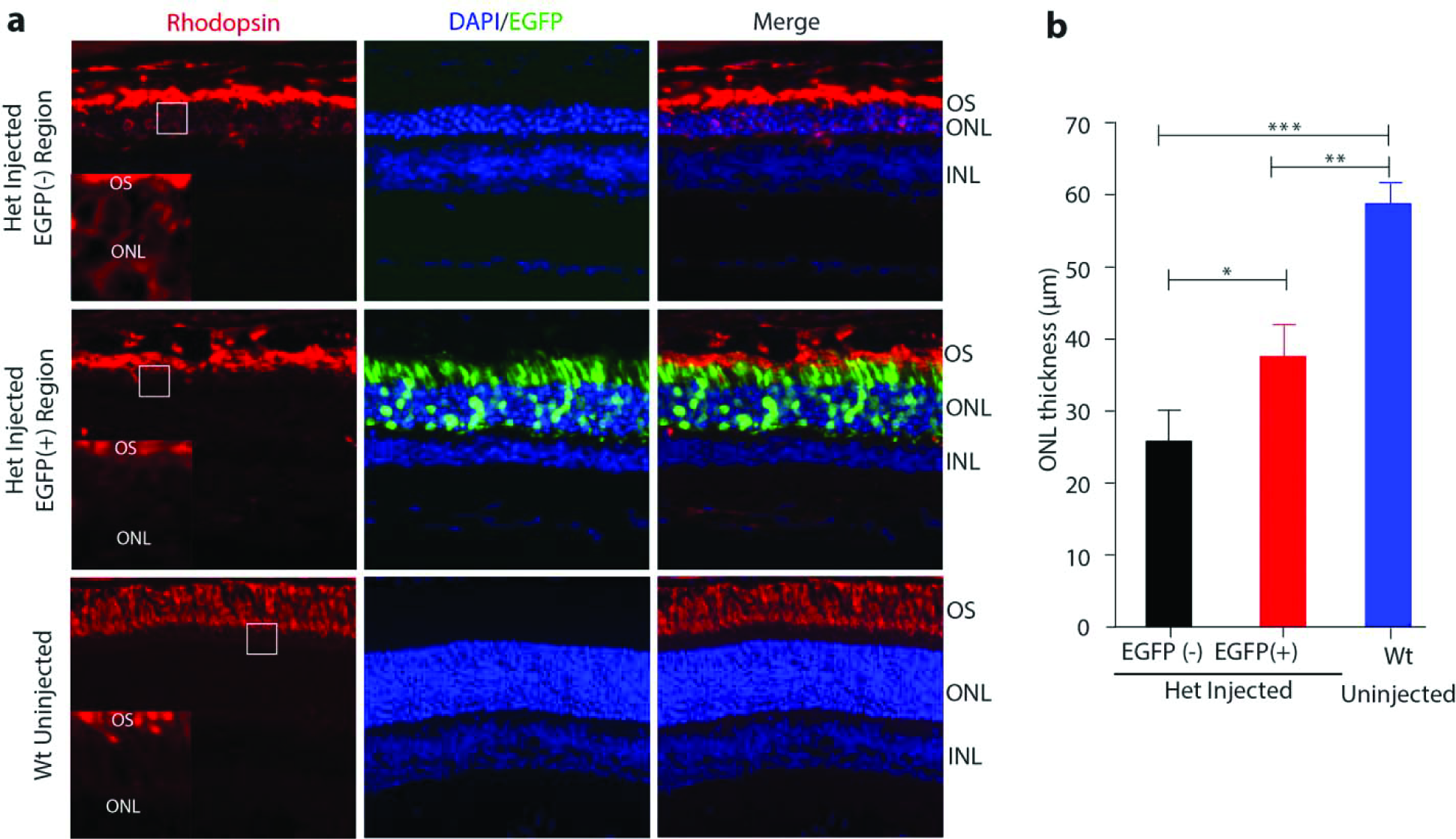
Photoreceptor cell preservation after gene editing with tru-sgRNA2 and SpCas9-VRQR in P23H heterozygous mice. **(a)** Retinal images from the treated region (EGFP+) and adjacent untreated region (EGFP-) in the P23H het mice at 5 weeks post-injection. Wild-type uninjected retina was used as control (bottom). Rhodopsin expression is detected by rhodopsin antibody (red), the EGFP signal indicates transfection (green), and nuclei were counterstained with Hoechst (blue). OS: outer segment, ONL: outer nuclear layer, INL: inner nuclear layer. Insets at the bottom right corner of each Rhodopsin panel were high magnification image of the white boxes. **(b)** Comparison of ONL thickness of the treated (EGFP+) area to its adjacent untreated (EGFP-) central area and uninjected wild-type controls (n=3). * *P* < 0.05; ** *P* < 0.01; *** *P* < 0.001 (Student’s T-test, n=3).

### Evaluation of off-target activity

To evaluate whether tru-sgRNA2 together with SpCas9-VRQR would cause any unwanted off-target cleavage in the mouse genome, we utilized targeted deep sequencing to assess off-target cutting at the top 10 potential tru-sgRNA2 binding sites out of 2,074 identified using CCTop (listed in *Supplementary Table 3*).^35^ NGS analysis showed no detectable off-target activity for 9 out of 10 candidate sites **(Fig. 5)**, which harbor between 2 and 4 mismatches compared the on-target P23H sequence **(Supplementary Table 3)**. Low off-target activity (3.12 ± 2.28%) was detected at an off-target site, which harbors a 1bp mismatch at position -11 from the PAM, in the region of a predicted miRNA, Gm27760, **(Fig. 5)**.

**Figure 5.**
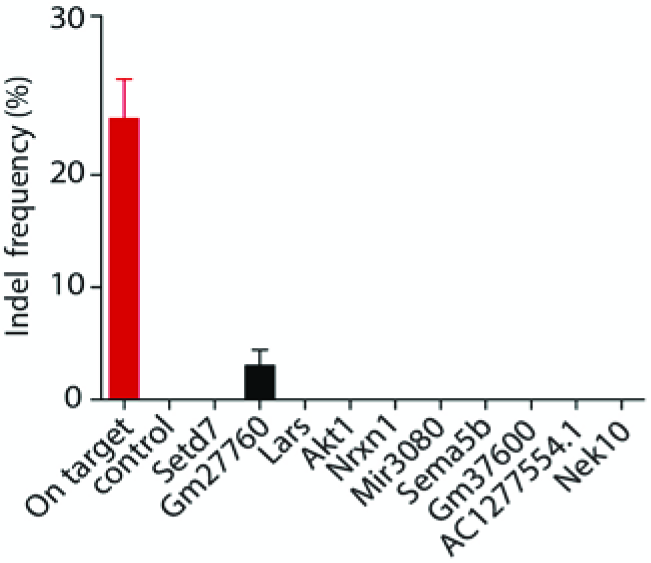
Off-target evaluation of tru-sgRNA2 with SpCas9-VRQR *in vivo*. Off-target analysis was conducted at the *in silico* predicted potential top 10 off-target sites. Injected retinas were collected and analyzed by PCR followed by targeted deep sequencing 7 days post-injection. Targeting activities on the P23H site (on-target) and 10 off-target sites were determined by NGS reads of PCR amplicons. Data is represented as mean ± SD (n=3). Control: uninjected mice.

## Discussion

Allele-specific CRISPR/Cas9 genome editing provides a promising means to treat dominantly inherited diseases. Here, we demonstrated that an *in vivo* allele-specific genome editing approach based on sequence differences in the spacer region of the sgRNA can selectively inactivate a single-base missense dominant allele at its native locus in the *RHO*-P23H knockin mouse model of adRP. By allowing for targeting of mutations that occur outside of PAM sequences, this approach significantly broadens the potential application of allele-specific CRISPR/Cas9 genome editing for the treatment of dominant genetic diseases.

We accomplished allele-specific targeting using a truncated sgRNA along with an SpCas9 variant designed to target an alternative PAM sequence. Although the targeting specificity of CRISPR/Cas9 is largely determined by the 10-12bp PAM-proximal seed sequences of sgRNAs,^14, 36-38^ it is notable that single mismatches within the seed sequence, even at the intended break point as in the case of sgRNA2, may not provide sufficient allele-specific targeting. This indicates that this spacer-mediated allele-specific knockout strategy works in a site-dependent manner. Thus, empirical evaluation of each individual allele-specific knockout approach is required.

It is particularly encouraging that, despite the inability of the 20-nt sgRNA2 to exhibit allele-specificity, a truncated 17-nt version of sgRNA2 paired with VQR version of SpCas9 variant provided exclusive allele-specificity in this study. It is also of interest that, in comparison to the VQR version of SpCas9 variant, the enhanced version of SpCas9-VRQR nuclease provided an approximately two-fold increase in on-target activity on the mutant allele, albeit with an increase in the targeting of the wild-type allele from 0% to 1.3 ± 0.3%. Notably, this low level disruption of the wild-type rhodopsin allele did not abrogate the observed therapeutic benefit, as evidenced by the significant increase of the relative levels of wild-type vs. mutant *Rho* mRNA in the treated cells **(Fig. 3)**. This finding is consistent with previous observation that an increased ratio of normal vs. mutant protein may be therapeutically beneficial for dominant diseases caused via a dominant negative mechanism.^39^ Additional studies will be needed to seek for the appropriate balance between cutting efficiency of the mutant allele and the level of modification of the normal allele.

It is worth mentioning that the allele-specific CRISPR/Cas9 gene knockout approach can be designed not only to target the mutation alone, but can also be broadened to common or rare SNPs that are in cis with the mutation. For example, the common SNPs that represent a given haplotype can all be used for allele-specific knockout of mutant alleles caused by different mutations,^10^ which could make the allele-specific therapeutic strategy more generic and cost effective. Moreover, as the expansion of the CRISPR/Cas toolbox continues, it may be possible to achieve highly efficient and allele-specific knockout of most, if not all, human dominant alleles via a similar allele-specific editing approach.

The potential off-target effects at unintended sites raises safety concerns for translating CRISPR/Cas9-based genome editing therapies to humans. In this proof-of-concept study, we performed off-target analysis for 10 *in silico* predicated sites and found a low-level off-target activity in a predicted miRNA Gm27760 in the mouse genome. No information regarding the function of this predicted miRNA is available to date. Since the tru-sgRNA2 cannot be translated to human use due to sequence mismatches between species, we elected not to study the effect of off-target activity at the Gm27760 site further. Rather, we anticipate more comprehensive studies of potential off-target effects associated with the development of a human specific P23H Cas9/sgRNA system, which we believe is warranted based on the results of this and other studies.^20, 40^ Furthermore, multiple means with the potential to improve targeting specificity, such as truncated gRNAs,^16^ Cas9 nickases,^41, 42^ and the more recently engineered Cas9s (eSpCas9,^43^ SpCas9-HF1,^32^ and HypaCas9^15^), can be explored to minimize undesired off-target activity.

Our study demonstrates that the allele-specific knockout of the mutant P23H allele in mouse photoreceptor cells results in a significant reduction of the mutant *Rho*-P23H mRNA in the transfected cells, which in turn leads to local preservation of photoreceptor cells in the treated regions of the *Rho*-P23H retinas. Two recent studies have reported successful CRISPR/Cas9-mediated cleavage of a human *RHO*-P23H allele in mouse and pig transgenic models, with similar phenotypic rescue observed in the *hRHO*-P23H transgenic mice.^20, 40^ Although encouraging, the disruption of the human *RHO*-P23H transgene was achieved by exploiting the sequence differences between the human transgene and the endogenous allele in the host animals.^20, 40^ Thus, further investigation is required to thoroughly validate the allele-specificity of the human *RHO* Cas9/sgRNA systems, including the SaCas9/sgRNA used in previous studies^20, 40^ and the corresponding VQR-SpCas9/tru-sgRNA2 system used in this study, in a heterozygous P23H human cell line or a humanized knock-in animal model harboring both a human P23H and a normal *hRHO* gene. It should also be noted that the therapeutic efficacy in our study was restricted by the delivery approach using plasmid DNA and *in vivo* electroporation. We are hopeful that other more relevant and effective *in vivo* delivery approaches, such as the use of dual adeno-associated virus (AAV) vectors encoding the Cas9 and sgRNAs separately, will result in treatment of more photoreceptor cells. This dual-vector approach has been successfully applied to the delivery of Cas9/sgRNA-mediated genome editing components into the mouse retina.^44-47^

## Conclusion

Our proof-of-concept study demonstrates that an allele-specific CRIPSR/Cas9 editing approach reduces photoreceptor cell death in *Rho*-P23H knockin mice by selectively targeting the endogenous mutant P23H allele. Our study also provides evidence that spacer-mediated allele-specific genome editing approaches are feasible for the treatment of dominant genetic diseases.

## Acknowledgements

This study is supported by grants from the Fighting Blindness Foundation (Q.L.); Research to Prevent Blindness Foundation (Q.L.); the National Eye Institute RO1 EY012910 (E.A.P., Q.L.) and P30 EY014104 (MEEI core support); Banting (Natural Sciences and Engineering Research Council of Canada) (B.P.K.); a Charles A. King Trust Postdoctoral Fellowship (B.P.K.); NIH R35 GM118158 (B.P.K., M.S.P., and J.K.J.); and a Desmond and Ann Heathwood MGH Research Scholar Award (J.K.J.).

## Authorship Confirmation Statement

Q.L. conceived the study. B.P.K. contributed to experimental design; B.P.K. and Q.L. designed the Cas9 and sgRNAs constructs; P.L., B.P.K. and M.S.P. generated plasmid constructs; P.L. and Q.L. designed and performed *in vivo* experiments with technical help from M.Y.L. and S.H.G.; P.L. and M.Y.L. executed *in vitro* experiments; P.L., D.N., and Q.L. analyzed the deep sequencing data; P.L. and Q.L. performed data analysis; E.A.P and J.K.J contributed to study conception and interpretation; P.L. and Q.L. drafted the manuscript; B.P.K., S.H.G., E.A.P., J.K.J. and Q.L. revised the manuscript. Q.L. supervised the overall project, and performed the final manuscript preparation. All authors reviewed and approved the manuscript prior to submission. The manuscript has been submitted solely to this journal and is not published, in press, or submitted elsewhere.

